# Data-driven lifespan transitions: cortical morphometry and intrinsic differences across network scales

**DOI:** 10.64898/2026.04.24.715834

**Authors:** Rose Cuthbertson, Oliver Edward Hancock, Vesna Vuksanovic

## Abstract

Age-related changes in cortical morphology are known to be nonlinear and feature-specific. However, age is often modelled using arbitrary bins or treated as a continuous variable. At the same time, structural covariance networks (SCN) have demonstrated that different morphometric features exhibit intrinsically distinct patterns of network organisation. Here, we integrate these approaches by introducing a data-driven framework to identify lifespan transitions in cortical morphometry and relate them to differences in SCN organisation. Using bootstrap-stabilised decision tree regression, we identify robust age partitions for multiple cortical morphometric features (surface area, its thickness and folding), revealing distinct feature-specific ageing regimes across lifespan (18 – 94 years of age). Our results show that features exhibiting similar lifespan transition profiles also demonstrate similar community-level SCN organisation, while features with divergent age trajectories show different network organisation. These findings demonstrate that lifespan transitions in cortical morphology are intrinsically linked to feature-specific network architecture, supporting the view that morphometric features capture distinct biological processes. Our results highlight the importance of data-driven lifespan modelling and reinforce the need to treat morphometric features as non-interchangeable when constructing network-based models of brain structure.

## Introduction

Cortical morphology changes continuously across the lifespan. However, emerging evidence suggests that different morphometric characteristics, usually estimated from *in vivo* magnetic resonance images (MRI), follow distinct age-related trajectories (15, 16, 24, 37). A fundamental understanding of the reorganisation of cortical morphometry across the lifespan should therefore address the transition points of such reorganisation. In addition, measures of cortical morphometry such as cortical thickness, surface area and curvature each reflect partially independent biological processes (6, 7, 30, 37) and their temporal characteristics. Thus, identifying their relationship with age related transitions will teach us about brain ageing and how biological processes unfold across the lifespan.

Most lifespan study approaches are based on predefined age bins or parametric assumptions about the relationship between age and the investigated outcome. However, while informative, these approaches may conceal feature-specific patterns of morphometric changes across the lifespan. Although changes in cortical thickness and volume have been extensively studied in ageing (12, 15), studies about cortical regional folding and geometry across the adult lifespan are lacking. Biologically, measures derived from cortical curvature, including Gaussian curvature, capture local geometric properties of the folded cortical surface. These measures are thought to reflect long-term mechanical and cytoarchitectonic constraints on cortical organisation (3, 4). As such, geometry-based models of cortical ageing predict that progressive, spatially heterogeneous tissue loss can lead to alterations in local curvature without changes to global folding topology (3–5). However, empirical studies in curvature-based ageing remain under-represented. Curvature measures complement information estimated from region-averaged cortical thickness of the highly convoluted cortical structure – they reflect how thickness changes are geometrically expressed across folds (11). Moreover, it would be interesting to know how age-related changes in cortical regional geometry relate to large-scale patterns of structural covariance organisation. Given the emerging evidence that suggests highly flexible topology of these networks folding-related covariance networks exhibit high flexibility across lifespan stages, suggesting that geometric features may act as late-life organisational markers of cortical ageing rather than following smooth, continuous trajectories.

Another, complementary perspective on morphological changes in the cortex is to look at its organisation through structural covariance networks (SCN), which quantify large-scale patterns of coordinated anatomical variation across individuals (10, 40). These networks consistently reveal modular and hierarchical structure across clinical conditions (20, 36, 42). However, it is not known whether features that age similarly also exhibit similar covariance organisation. Establishing such a link would offer insight into whether lifespan changes in cortical morphology relate to intrinsic architectural properties of the cortex, rather than reflecting independent or purely local effects. We quantify similarity of network community structure across age regimes using adjusted Rand index, a well-established measure of network reorganisation and similarity between networks, associated with each morphometric feature.

Here, we introduce a data-driven framework that identifies lifespan transitions in multiple morphometric features using bootstrap-stabilised decision-tree regression, thereby allowing transition points to emerge directly from the data, revealing feature-specific ageing profiles. We then relate these profiles to the intrinsic organisation of sex-corrected structural covariance networks, assessed via community structure stability across data-driven age bins, and assess which regional features are most informative for age prediction using an interpretable machine-learning framework. Together, these approaches allow us to investigate whether morphometric features that undergo similar lifespan transitions also share common patterns of large-scale cortical organisation.

## Methods

### Imaging Dat

Anatomical T1-weighted MRI scans from healthy individuals aged 18–94 years from four publicly available datasets (OASIS-1, OASIS-2, NKI, and IEEE) were combined to obtain a multi-site dataset of age-related morphometric features. A demographic breakdown of the combined dataset and individual cohorts’ demographics is provided in Table 1. Cortical measurements were extracted using the Destrieux atlas parcellation, which provides gyral- and sulcal-level anatomical resolution (9), and divides the cortical surface into 148 regions of interest. Image processing was performed on the Super Computing Wales high-performance cluster within FreeSurfer pipelines (v.6.0) using *recon_all* function and its default setup (2). Cortical morphometric measures were derived from FreeSurfer output using a standard in-house Matlab script (41), yielding region-wise estimates of surface area, grey matter volume, cortical thickness, folding index, and curvature-based measures (Gaussian and mean curvatures).

**Table 1.** Demographic characteristics of the datasets.

| Dataset | Age range (years) | Total | Males | Females |
| --- | --- | --- | --- | --- |
| Total | 18–94 | 1009 | 456 | 553 |
| IEEE | 45–86 | 340 | 159 | 181 |
| NKI | 18–83 | 154 | 95 | 59 |
| OASIS-1 | 18–94 | 379 | 147 | 232 |
| OASIS-2 | 60–93 | 136 | 55 | 81 |

Because data were acquired across different scanners and acquisition protocols, site-related technical variability unrelated to biological effects was corrected through multi-site harmonisation performed using the neuroHarmonize Python package (14, 19, 29). This method is based on the ComBat harmonisation framework and incorporates generalized additive models (GAMs) to model and preserve nonlinear covariate effects, including age. To evaluate the effectiveness of the harmonisation procedure, one-way ANOVA was performed to assess site-related differences in morphometric measures across datasets. Prior to harmonisation, the mean F-statistic across regions was 115.08, indicating substantial site effects. Following harmonisation, this value was reduced to 8.67. Residual differences are expected, given the non-overlapping age ranges across datasets, which may introduce biologically meaningful variation in the harmonised morphometric measures.

### Data-Driven Lifespan Partitioning and Feature Importance

Decision tree regression with age as the predictor and each morphometric feature as the outcome variable, was used to identify age ranges associated with age-related transitions in cortical morphometry. Data were split into training and test sets, with 807 cases used for training and 202 for testing. To ensure robustness, the procedure included bootstrap resampling: For a given feature, shallow decision trees were fitted to bootstrap resamples of the data (on 1,000–5,000 iterations), and the resulting age-dependent splits were aligned across iterations using their mean and confidence intervals. This approach provided stable, data-driven age partitions for each feature. The partitions’ split points were then converted into adjacent age ranges spanning the full lifespan. To evaluate the validity of the resulting data-driven partitions, feature values were grouped according to the derived age ranges and compared using one-way ANOVA (F- and associated p-values are reported for between-group separation). The F-values show how strongly that feature separates the data-driven age groups. This procedure was applied independently to each morphometric feature to obtain feature-specific lifespan partitions and their statistical characterisations (see Table S1 for partitions’ demographic characteristics).

Hierarchical clustering, on feature-specific vectors summarising the locations of data-driven age partitions, was used to compare lifespan transition profiles across morphometric features. Clustering was carried out using Ward’s linkage criterion, which groups features based on similarity in their lifespan transition structure while minimising within-cluster variance. The resulting dendrogram was used to determine an ordering of features that reflects shared patterns of lifespan transitions. This ordering was then used to visualise feature-specific age ranges as horizontal bar plots spanning the full lifespan, allowing systematic similarities and differences in the timing and extent of age regimes to be compared across features in a unified framework.

In addition, an Explainable Boosting Machine (EBM) – an interpretable additive model that represents predictions as a sum of feature-specific effects – was used on the test dataset to quantify feature importance in terms of their age-dependent contribution profiles (25). EBM was chosen because it provides transparent, non-linear feature contributions at the individual level, allowing age-dependent effects to be characterised without imposing parametric assumptions. For each morphometric feature, subject-level contributions were averaged across age and smoothed to obtain lifespan contribution curves, with uncertainty estimated via bootstrap resampling. Global feature importance was quantified as the mean absolute contribution across subjects. Age contribution trajectories were computed from local explanations of the final model on the test set. The local explanations of the model break a prediction down into its additive term contributions. Using the test set to model the contribution of terms across age uncovers within which age ranges are specific regions/features the most informative for predicting age. For each term, a locally estimated scatterplot smoothing (LOESS) model was fitted relating contribution to age. LOESS was chosen as an easy to use, non-parametric, non-linear model that is robust to outliers (8). The trajectory was then derived by using each term’s LOESS model to make a prediction of the contribution across 100 equidistant points between the minimum and maximum age. To quantify uncertainty in the estimated trajectories, 1000 iterations of bootstrap resampling were performed. Within each bootstrap, a LOESS model was fit and predictions made across the same 100 points. This method of confidence interval estimation was chosen due to its ability to handle sparsity in the underlying age distribution.

### Cortical Similarity Network Construction and Analysis

Structural covariance networks were constructed at the group level by computing partial correlations between regional morphometric measures across participants within each of the four data-driven lifespan groups, controlling for sex, following established approaches (42). This procedure removes covariance due to sex differences while preserving intrinsic inter-regional relationships. For each morphometric feature (8 in total), four sex-corrected SCNs were estimated using bootstrap resampling of 100 participants per lifespan group. This sample size was chosen to match the smallest group in the whole cohort (see Supplementary Table 1.) and was applied consistently across all age groups to ensure comparability of network estimates and to mitigate potential biases arising from unequal group sizes (42).

### Community Structure Estimation and Similarity Analysis

Community structure was estimated for each feature-specific SCN using modularity maximisation applied to signed networks (32), similar to our previous work (42). To provide the robustness of community organisation estimates, 200 Louvain repetitions per age group for consensus were used. This approach allowed us to estimate feature-specific, intrinsic community organisation while accounting for sampling variability. Community structures were detected using the Brain Connectivity Toolbox (32) function *community_louvain* with the following inputs: *γ* = 1.0 and symmetric treatment of negative weights (^*′*^*negative*_*sym*^*′*^) (42).

Similarity between community structures was quantified using the adjusted Rand index (ARI), a chance-corrected measure of agreement between two partitions of the same node set (17). ARI assesses the consistency of pairwise node assignments to communities across partitions, correcting for agreement expected by chance. Values range from -1 to 1, with higher values indicating greater similarity between community structures. ARI was calculated between adjacent lifespan splits on feature-specific, sex-corrected SCNs. This way eight 4*×*4 similarity matrices were obtained, whit entries representing estimated pair-wise ARIs.

## Results

### Feature-Specific Lifespan Transitions

Figure 1 shows data-driven lifespan partitions revealing distinct, feature-specific ageing regimes. Feature importance in driving four data-driven age-related splits across the lifespan was quantified by the F-test value: Regional thickness average and surface area showed earlier and more pronounced age splits, while curvature-related measures demonstrated less pronounced (see corresponding F-values) or later-occurring transitions. We highlighted contributions of cortical surface area and its average thickness, rather than regional gray volume because volume is a combined measure of the two characteristics and its importance clearly arises from these combined effects. The same information is summarised in Figure2, highlighting feature-specific similarities and differences in the timing of cortical transitions.

**Fig. 1.**
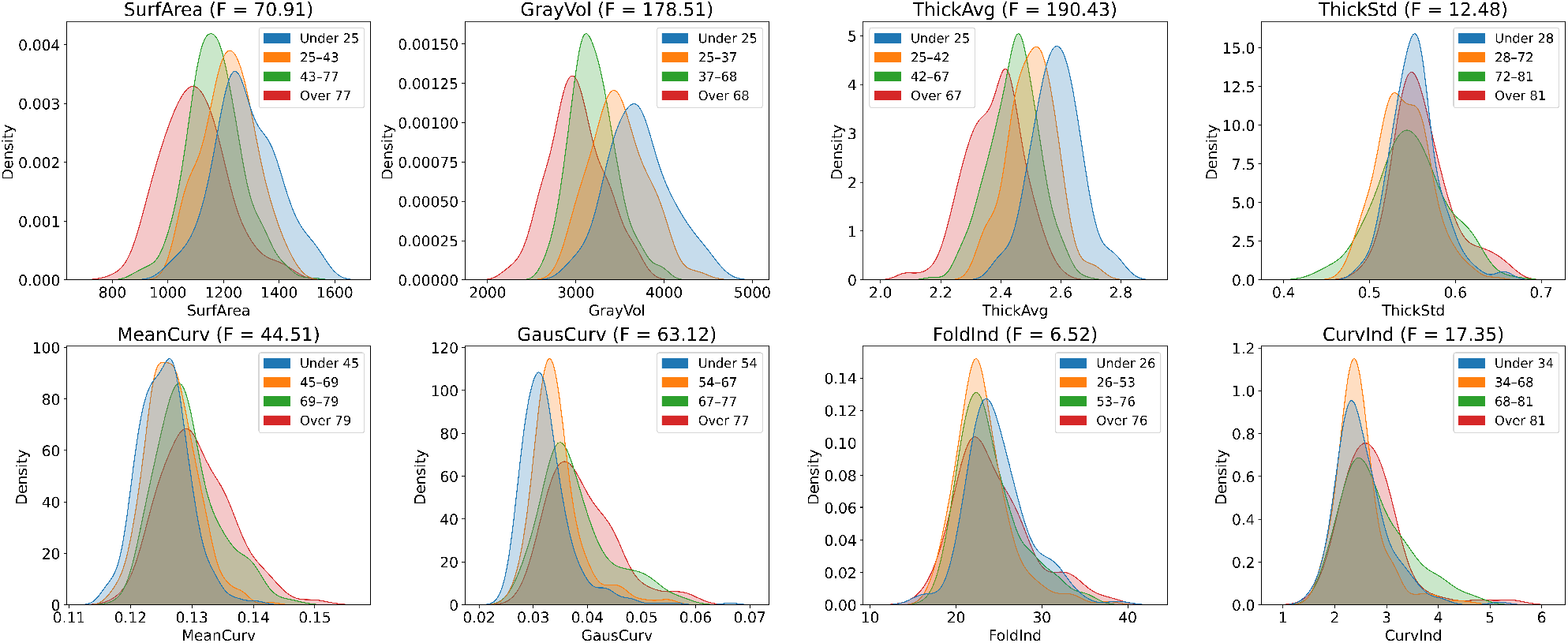
Data-driven lifespan partitions across cortical morphometric features. Kernel density estimates of morphometric feature distributions are shown across age groups defined by bootstrap-stabilised decision tree regression. Age partitions differ systematically across features, indicating distinct lifespan transition profiles. F-statistics quantify between-partition separation for each feature.

**Fig. 2.**
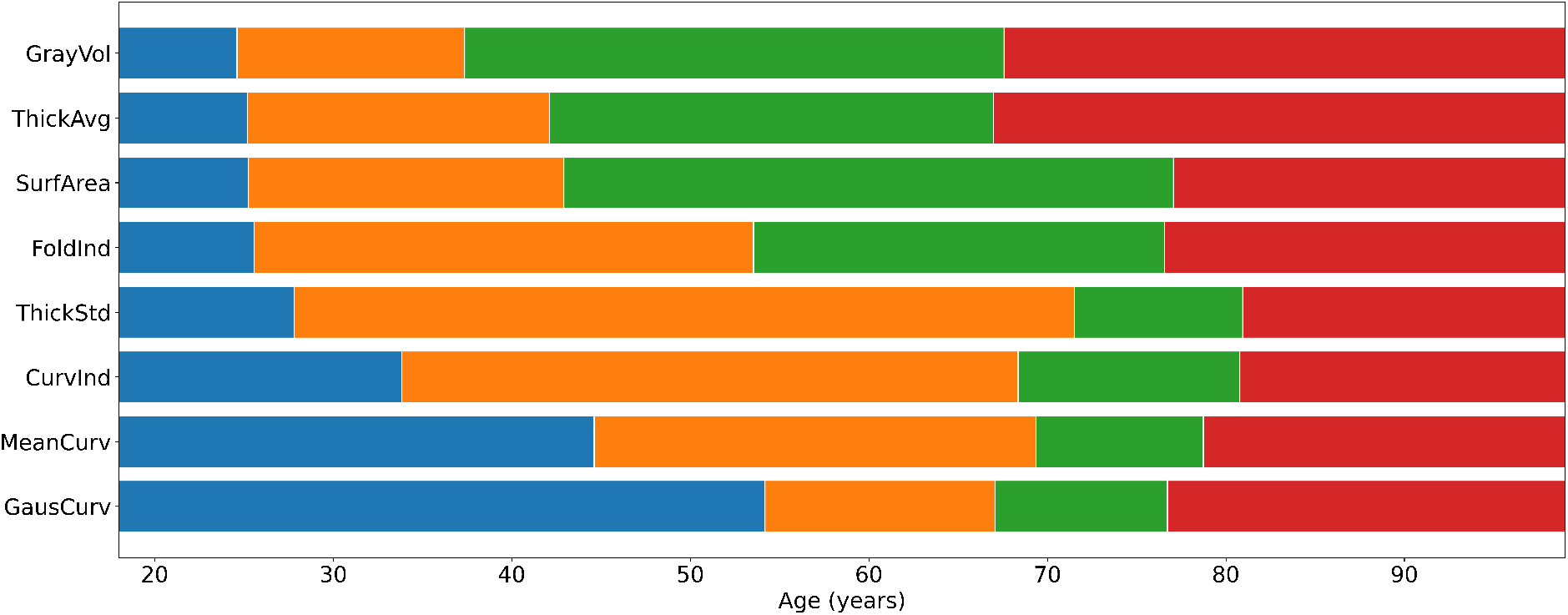
Cross-feature data-driven lifespan transition profiles. Lifespan partitions derived from the bootstrap-stabilised decision tree analysis shown in Figure 1 are summarised for each morphometric feature. Coloured segments indicate age regimes defined by the mean split locations, with features ordered by hierarchical clustering of split positions. This synthesis highlights systematic similarities and differences in the timing of lifespan transitions across morphometric features.

For each feature, coloured segments indicate the age regimes defined by its mean split locations, providing a compact view of its inferred lifespan trajectory (colour codes match those in Fig. 1). The features were ordered by hierarchical clustering, which reveals that some features, such as cortical regional thickness or volume, show earlier transition points (below 30 years of age). In contrast, curvature-related features, e.g., mean and Gaussian curvatures, display later transitions, indicating more prolonged lifespan trajectories. The visual comparisons of these regimes highlights systematic similarities and differences across morphometric features, demonstrating that cortical morphology does not follow a single, uniform ageing pathway but instead exhibits distinct, feature-specific patterns of reorganisation. These findings indicate that morphometric features capture different aspects of cortical ageing rather than reflecting a single unified ageing process.

### Feature Importance

Table 2 summarises results of feature importance analysis, which quantifies the relative contribution of each region/feature to the lifespan splits for that feature. The 10 most important features were comprised primarily of thickness (n = 5) and curvature measures (n = 4), and one gray volume (for cuneus gyrus). Feature importance analysis showed that average cortical thickness was the most important main effect, with thickness of the calcarine sulcus emerging as the single most important feature (importance: 2.6). Five important features were located in the occipital, three in the frontal and two in the temporal lobe. Contribution trajectories of the three main features across the lifespan are shown in Fig. 3. The contribution trajectories reveal that the single most important feature, the average of thickness of the calcarine sulcus, has a consistent linear-like trend contribution to the age prediction spanning from -3.08 years at 18 to +3.37 years at 94, implicating it as an important biomarker for age across the lifespan. The mean curvature of the cuneus gyrus provides a slight negative contribution to the prediction up to ≈60 years, after which it provides no contribution at all until ≈75 years, and from there on provides a steep negative contribution that reaches -1.68 years at 94 years. Indicating that this main effect has the most discriminative power when predicting ages above ≈75. Gaussian curvature of the middle occipital gyrus shows a steady, small negative contribution across most of the lifespan. However, after around the age of ≈60 it then sharply increases up to +3.95 years of contribution at age 94.

**Table 2.** Ranked main features (effects) with corresponding importance values.

| Rank | Feature | Importance |
| --- | --- | --- |
| 1 | Calcarine Sulcus Thickness Avg | 2.6 |
| 2 | Cuneus Gyrus Mean Curvature | 1.9 |
| 3 | Occipital Middle Gyrus Gaussian Curvature | 1.9 |
| 4 | Frontal Superior Gyrus Thickness Avg | 1.5 |
| 5 | Temporal Inferior Sulcus Gaussian Curvature | 1.5 |
| 6 | Frontal Middle Gyrus Thickness Avg | 1.3 |
| 7 | Frontal Superior Gyrus Gaussian Curvature | 1.2 |
| 8 | Temporal Superior Sulcus Thickness Avg | 1.2 |
| 9 | Cuneus Gyrus Grey Matter Volume | 1.1 |
| 10 | Occipital Middle Gyrus Thickness Avg | 0.9 |

**Fig. 3.**
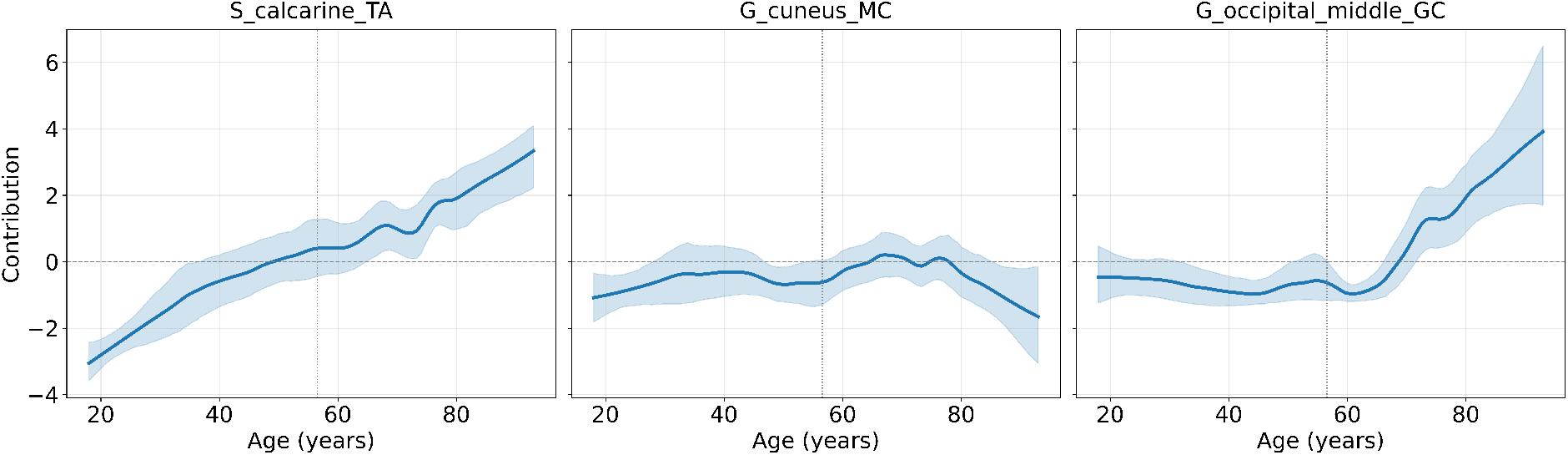
Age contribution trajectories of the top three main-effects. Contribution trajectories of the top three main-effects with 95% confidence intervals. The vertical dashed line shows the predicted mean that the model uses as the baseline estimate from which terms interact with additively (56.6)

### Linking Lifespan Transitions to Structural Network Organisation

Figure 4 exemplifies similarity between community structures across age groups quantified using the adjusted Rand index. ARI values vary across age transitions: small ARIs indicate low similarity, i.e., high reorganisation of covariance communities at corresponding lifespan transitions; higher ARIs indicate relative preservation of network topology across corresponding transitions. Because SCNs were estimated within age-corrected groups and controlled for sex, these differences reflect changes in structural covariance organisation associated with lifespan stage, rather than global sex effects.

**Fig. 4.**
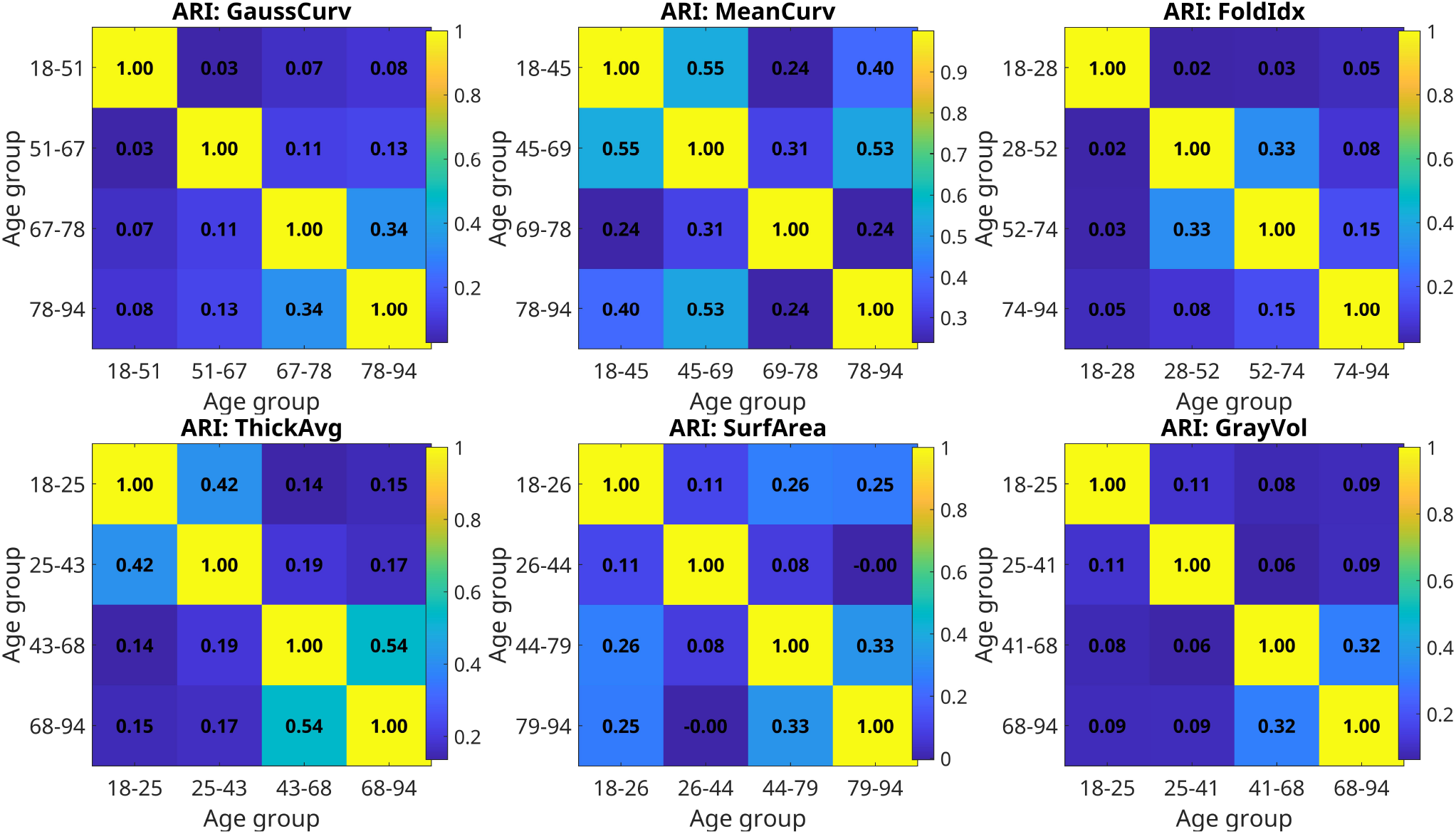
Similarity between community structures across data-driven age groups. Similarity between community partitions of structural covariance networks across the four lifespan groups (see Fig. 1) was quantified using the adjusted Rand index (ARI), yielding a 4×4 matrix for each morphometric feature. For each feature, SCNs were constructed separately within age groups using partial correlations controlling for sex, and community structure was estimated using signed modularity maximisation. Results are shown for six representative cortical features, illustrating feature-specific patterns of network reorganisation across the lifespan.

Here, a 4*×*4 matrix represents ARIs calculated on adjacent age-bins based on a single realisation of these networks. Distributions of ARIs across feature-level transitions are shown in Supplementary Information section **??**. Distributions drawn from 100 bootstraps (each) were calculated on sex-corrected SCNs across (i) adjacent age-bin-transitions within data-driven age bins (see Figs. S2 and S1), and (ii) across lifespan (see Fig. S3). Mean ARIs across bootstrapped samples between adjacent age bins for youngest and oldest bins are shown in Fig. 5.

**Fig. 5.**
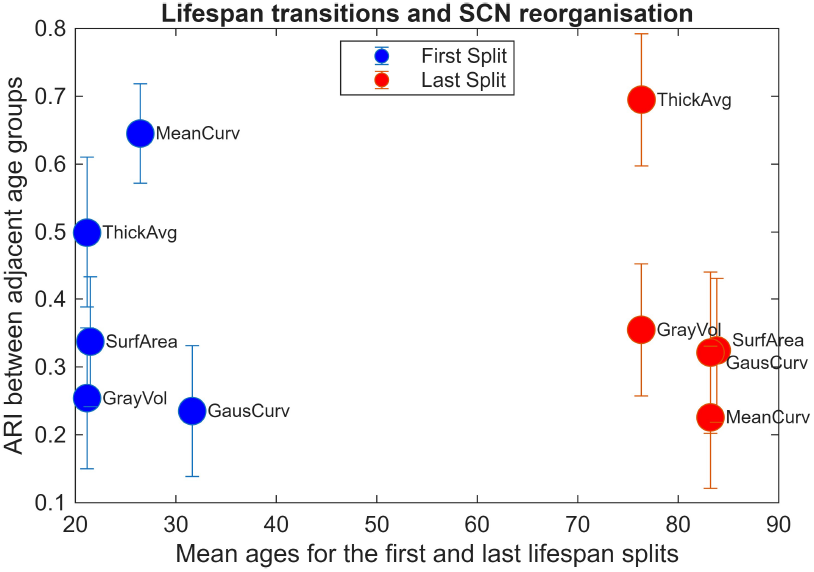
Feature-level relationship between lifespan complexity and structural covariance network reorganisation. Mean ARIs between first and second age groups were plotted against corresponding first lifespan split mean age. Abbreviation: Adjusted Rand index (ARI).

To understand linear contribution of age to SCN reorganisation across the lifespan, we calculated ARI on SCNs constructed on age-corrected vs age-included across the adult lifespan (see Fig. 6). Reorganisation of regional thickness, folding and Gaussian curvature show linear dependency with age, while gray volume, surface area and mean curvature show non-linear association with age. Similar can be concluded from the contribution trajectories of the three main features (see Fig. 3), which all display linear behaviour across the lifespan (thickness) or later-life years (mean and Gaussian curvatures).

**Fig. 6.**
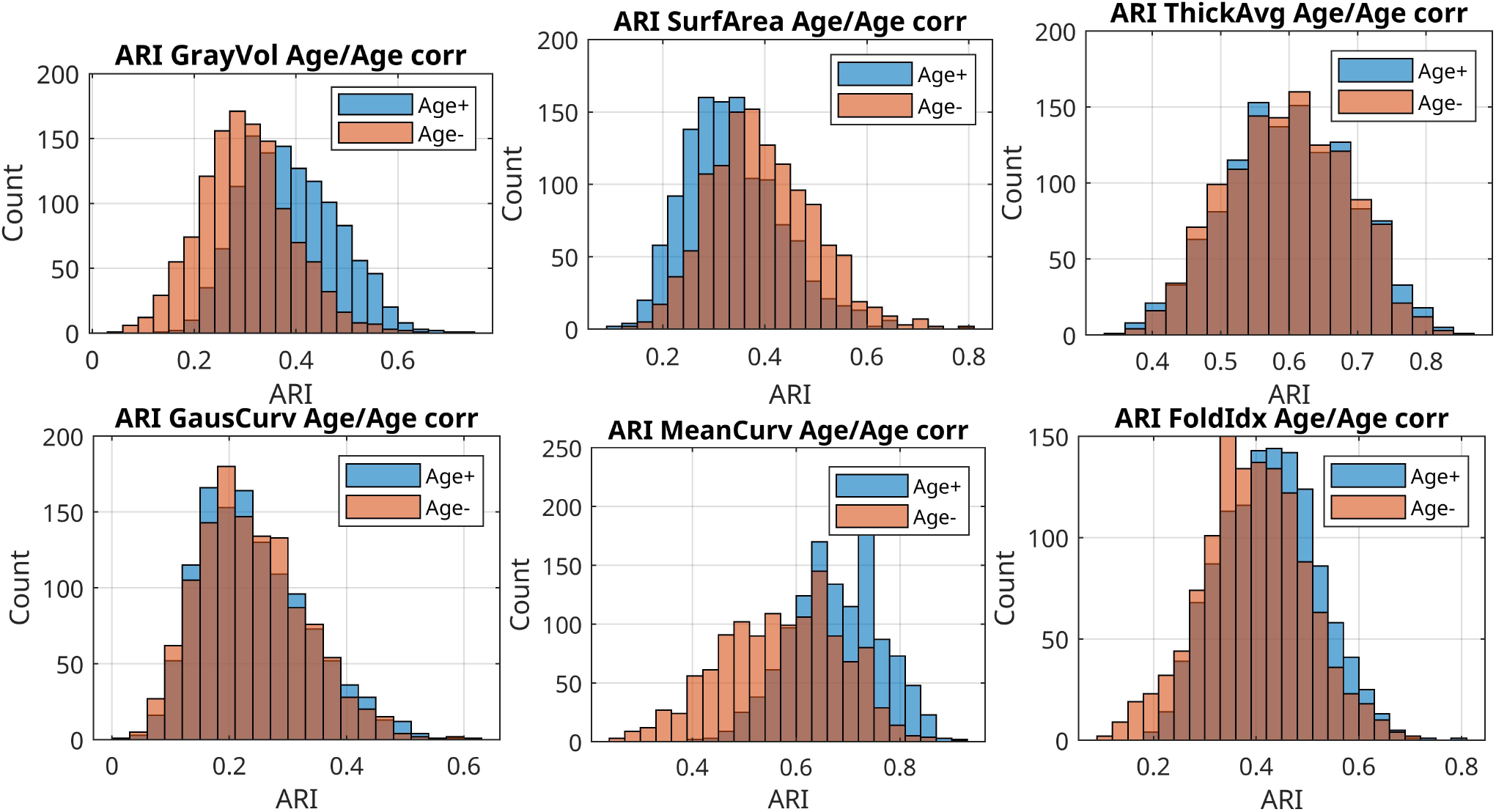
Feature-level lifespan structural covariance network reorganisation. Distribution of ARIs within lifespan age (18 – 94) calculated on SCNs corrected for age covariance (Age-) and not (Age+). Abbreviation: Adjusted Rand index (ARI); Structural covariance networks (SCNs).

## Discussion

In this study we combined data-driven lifespan partitioning with structural covariance network analysis in around 1000 participants between 18 and 94 years of age to demonstrate that age-related changes in cortical morphology are both strongly feature-specific and intrinsically linked to large-scale network (re)organisation. Our results show that different morphometric features exhibit distinct lifespan transition profiles, and that these transitions are reflected in systematic reorganisation of community structure within feature-specific SCNs. Together, these findings support the view that cortical morphometric features capture partially independent biological processes that unfold on different temporal and network scales.

First, the data-driven lifespan partitioning revealed clear differences in the timing and complexity of age-related transitions across morphometric features (Figs. 1 and 2). Cortical thickness, surface area, and grey matter volume showed earlier and more frequent transitions (see Fig. 2), whereas curvature-based measures exhibited later splits. Importantly, these feature-specific transitions emerged directly from the data, without reliance on predefined age bins or ageing models often used in ageing studies. As a result, the identified lifespan regimes capture intrinsic differences in how morphometric features evolve across the lifespan, highlighting that cortical-morphometry ageing is characterised by multiple, feature-dependent temporal profiles rather than a single, uniform trajectory. This pattern is consistent with prior work indicating that thickness-related measures are particularly sensitive to age-dependent processes such as early-age synaptic pruning and dendritic reorganisation (15, 33), or later-life-related neurodegeneration and tissue shrinkage(13, 21, 23, 31), while surface area and folding-related metrics may reflect more stable developmental and architectural properties of the cortex, shaped also by external, environmental rather than inherited factors (18, 38), which drive brain ageing patterns in later life.

Second, we identified which specific brain regions and morphometric features were most informative for predicting age, using an explainable machine learning model. The model identifies the contribution of regional features to (the model) performance for predicting age within the data-driven age bins. Thickness- and curvature-based measures contribute most strongly to the data-driven identification of age-related transitions, with particularly strong signal in frontal and occipital regions (see Table 2). Frontal regions are known to undergo marked structural change across the lifespan and are among the earliest to show age-related alterations in both healthy ageing and neurodegenerative conditions (1, 35). The dominance of thickness-related measures is consistent with prior evidence that laminar microstructural organisation is especially sensitive to age-related processes, including synaptic and dendritic reorganisation, whereas the contribution of curvature-based measures highlights the additional role of cortical geometry in shaping age-related variability (3, 4). As expected, the occipital lobe shows the strongest contribution, with the top three regions all belonging to this cortical lobe. Previous studies have shown that ageing does not tend to affect the occipital cortex at similar stage as e.g., frontal and temporal regions (1, 27). The occipital lobe is consistently noted among regions with the least age-associated structural (e.g., cortical thinning (34)) or functional (e.g., glucose uptake (43) or cognitive specialisation (28)) changes. Our results corroborate these findings suggesting that distinct biological processes – microstructural tissue properties and regional folding geometry –jointly contribute to cortical ageing, but do so with different regional importance and temporal profiles. As in earlier studies on heterogeneous atrophy patterns across the cortical surface in both healthy ageing and dementia (22, 39), these results underscore that age-related structural change is neither spatially uniform nor driven by a single morphometric mechanism.

Third, the observed lifespan transitions were not limited to regional morphometric changes but were accompanied by systematic differences in large-scale morphometric covariance organisation. Comparison of SCN community structure across the four data-driven age groups revealed marked variation in ARI values across features and transitions (see Fig. 4). Lower ARI values between specific adjacent age groups indicate reorganisation of network communities at corresponding lifespan stages, whereas higher ARI values suggest relative preservation of network topology across adjacent age ranges. Because SCNs were constructed within age-stratified groups and controlled for sex, these differences cannot be attributed to global age trends or sex effects. Instead, they point to intrinsic changes in how regional morphometric variations co-vary across the cortex at different stages of the lifespan. Features characterised by earlier or more complex lifespan transitions tended to show greater changes in community structure across age groups (Fig. 5), whereas features with later or more gradual transitions exhibited more stable network organisation. This relationship suggests that morphometric features vary not only in their temporal progression (ageing) but also how they are embedded within the network. Our findings align with prior work showing that the modular organisation of structural covariance networks depends critically on the morphometric features used to construct them, rather than reflecting a fixed property of cortical architecture (41). Extending this framework, we show that even feature-specific networks undergo systematic reorganisation across data-driven lifespan stages. Differences in community structure across age groups, quantified using the adjusted Rand index, indicate varying degrees of network reconfiguration, with lower ARI values reflecting greater reorganisation—or increased flexibility—of covariance communities across the lifespan. Features characterised by similar lifespan transition profiles tend to exhibit more preserved community organisation, whereas features with distinct transition timings show more pronounced modular reorganisation. Together, these results support the view that age-related cortical change is governed by multiple, feature-dependent organisational principles, and that ARI-based comparisons provide a concise measure of how structural covariance networks reorganise across lifespan stages.

Finally, we show that features with similar lifespan transition profiles tended to exhibit similar patterns of network reorganisation, whereas features with divergent ageing trajectories showed more distinct community structure changes (see Fig. 6). Across all morphometric features, age correction systematically alters lifespan SCN reorganisation, as reflected by shifts in ARI distributions. In most measures, age-corrected networks show higher or more stable ARI values, indicating increased consistency of network partitions across the lifespan. This suggests that a substantial component of apparent network reorganisation is driven by shared age-related covariance. Nonetheless, residual differences after correction highlight feature-specific, age-independent network dynamics. Cortical thickness shows little or no shift because age-related variance is already the dominant and relatively uniform signal in thickness across the cortex. Thickness declines with age in a highly global, monotonic, and spatially coherent manner, so correcting for age removes variance that is already largely shared across regions and subjects, without substantially altering the relative covariance structure that defines the SCNs. In contrast, measures such as surface area, curvature, or folding index capture more region-specific and developmentally heterogeneous processes, where age correction more strongly reshapes inter-regional covariance.

Our results suggest complex, *not* one-to-one, mapping between regional change in cortical morphometry and network organisation. We suggest that lifespan transitions represent points at which coordinated patterns of inter-regional variation reorganise, potentially reflecting shifts in the balance between local tissue properties and global cortical architecture. For example, Gaussian curvature is particularly sensitive to late-life geometric reorganisation of the cortical sheet (26). As a local folding metric, it captures subtle shape-changes arising from the interaction of the principal curvatures, making it responsive to sulcal sharpening, gyral thinning with preserved surface topology, and differential tissue loss across sulcal walls (2, 34). In this case, the observed increase in Gaussian curvature with age does not imply increased folding – it reflects the geometric consequences of ageing-related cortical thinning, which occurs non-uniformly across gyri and sulci, as well as sulcal widening and deepening due to volume loss. Although major folding patterns are established early in development, ageing-related cortical thinning and differential tissue loss across sulcal and gyral regions can lead to progressive changes in local curvature without altering overall surface topology. These processes can exaggerate local curvature values without altering overall folding patterns. Thus, while cortical thickness primarily reflects laminar tissue loss, Gaussian curvature captures how that loss is geometrically expressed across the folded cortex. The two measures are therefore complementary rather than contradictory, explaining why Gaussian curvature shows gradual age effects and a strong association with network reorganisation in the present study.

In conclusion, our results have important implications for network-based studies of brain structure and ageing. First, they demonstrate that morphometric features should not be treated as interchangeable proxies for cortical organisation. Different features capture distinct biological and network-level processes and characteristics, and merging across them may obscure differences in ageing patterns. Second, they highlight the limitations of conventional age modelling approaches that rely on predefined bins or linear parametric assumptions. Data-driven identification of lifespan transitions provides a framework to capture nonlinear ageing effects and to relate them to changes in network organisation.

### Limitations and Future Directions

Our analysis is cross-sectional, and although the data-driven partitioning reveals robust age-associated transitions, longitudinal data will be required to determine whether these transitions reflect within-individual change or cohort effects. Additionally, while multi-site harmonisation substantially reduced scanner-related variability, some residual site effects may remain. Future work could also extend our framework to include functional connectivity, microstructural characteristics, or molecular biomarkers to further elucidate the biological mechanisms underlying feature-specific ageing.

Despite these limitations, the present study provides a data-driven framework of how cortical morphometry and structural covariance organisation co-evolve across the human lifespan. By showing that lifespan transitions and network architecture are intrinsically linked in a feature-specific manner, our results highlight the value of large-scale, data-driven, multi-scale approaches in the study of brain ageing.

## Data and Code Availability

All analyses were implemented using MatLab (MathWorks, Natick, MA) and Python, and the source code and sample data can be obtained at the project’s OSF folder or from the corresponding author upon request.

## Authorship Contribution Statement

VV: Conceptualised the study and supervised the project, designed the analytical framework, interpreted results, and wrote the manuscript. RC: Performed formal analyses related to lifespan transitions. OEH: Performed structural covariance network analyses.

## Acknowledgment

We thank Venia Batziou for assistance with data preprocessing, project: DSR-1085-100.

## Supplementary Tables & Figures

**Table S1.** Demographic characteristics of the datasets.

| Feature | Age range (years)/N | Age range (years)/N | Age range (years)/N | Age range (years)/N |
| --- | --- | --- | --- | --- |
| GrayVol | < 25 / 146 | 25–37 / 146 | 37–68 / 405 | > 68 / 372 |
| ThickAvg | < 25 / 146 | 25–42 / 99 | 42–67 / 395 | > 67 / 372 |
| SurfArea | < 25 / 146 | 25–43 / 96 | 43–77 / 618 | > 77 / 124 |
| FoldInd | < 26 / 176 | 26–53 / 180 | 53–76 / 449 | > 76 / 257 |
| MeanCurv | < 45 / 260 | 45–69 / 401 | 69–79 / 212 | > 79 / 139 |
| GausCurv | < 51 / 345 | 51–67 / 203 | 67–77 / 265 | > 77 / 139 |

**Fig. S1.**
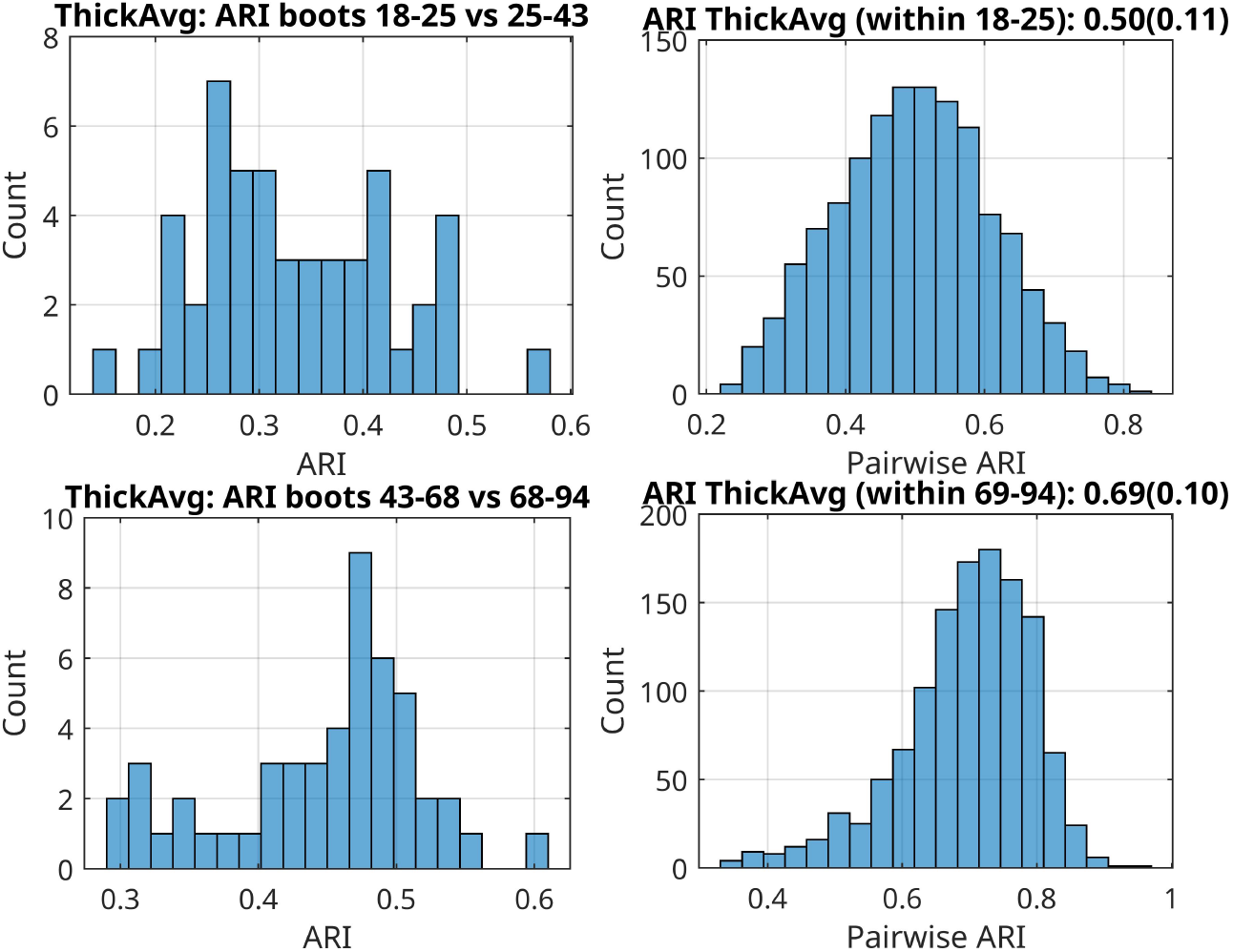
Thickness Average-level within- and between-age-bin structural covariance network reorganisation. Left panels (top and bottom): Distribution of ARIs within data-driven age-bins. Middle Panels: Distribution of ARIs between data-driven age-bins. Right panels: ARIs distribution across lifespan – sex-corrected (top) and age and sex corrected (bottom). Abbreviation: Adjusted Rand index (ARI).

**Fig. S2.**
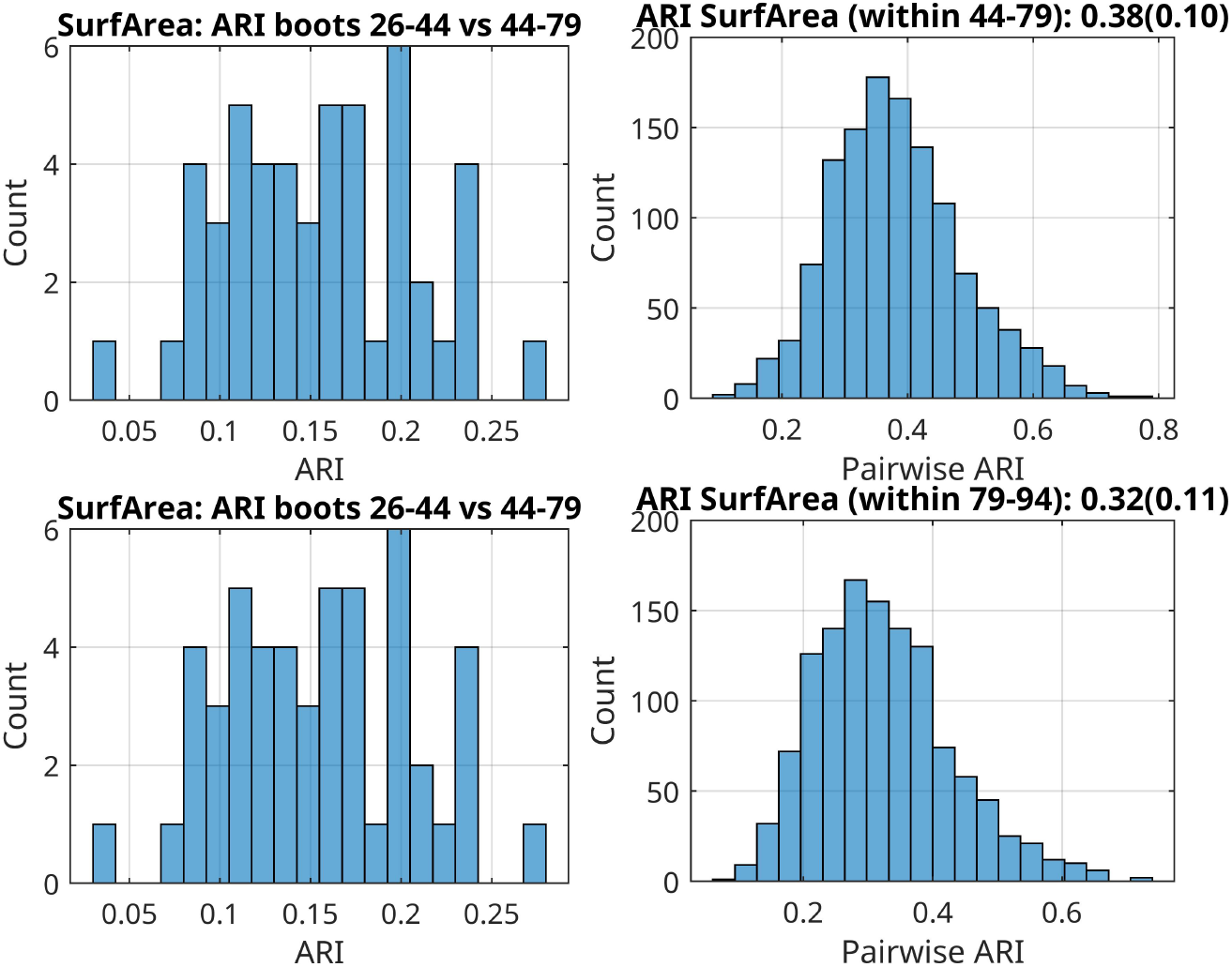
Surface Area feature-level within- and between-age-bin structural covariance network reorganisation. Left panels (top and bottom): Distribution of ARIs within data-driven age-bins. Middle Panels: Distribution of ARIs between data-driven age-bins. Right panels: ARIs distribution across lifespan – sex-corrected (top) and age and sex corrected (bottom). Abbreviation: Adjusted Rand index (ARI).

**Fig. S3.**
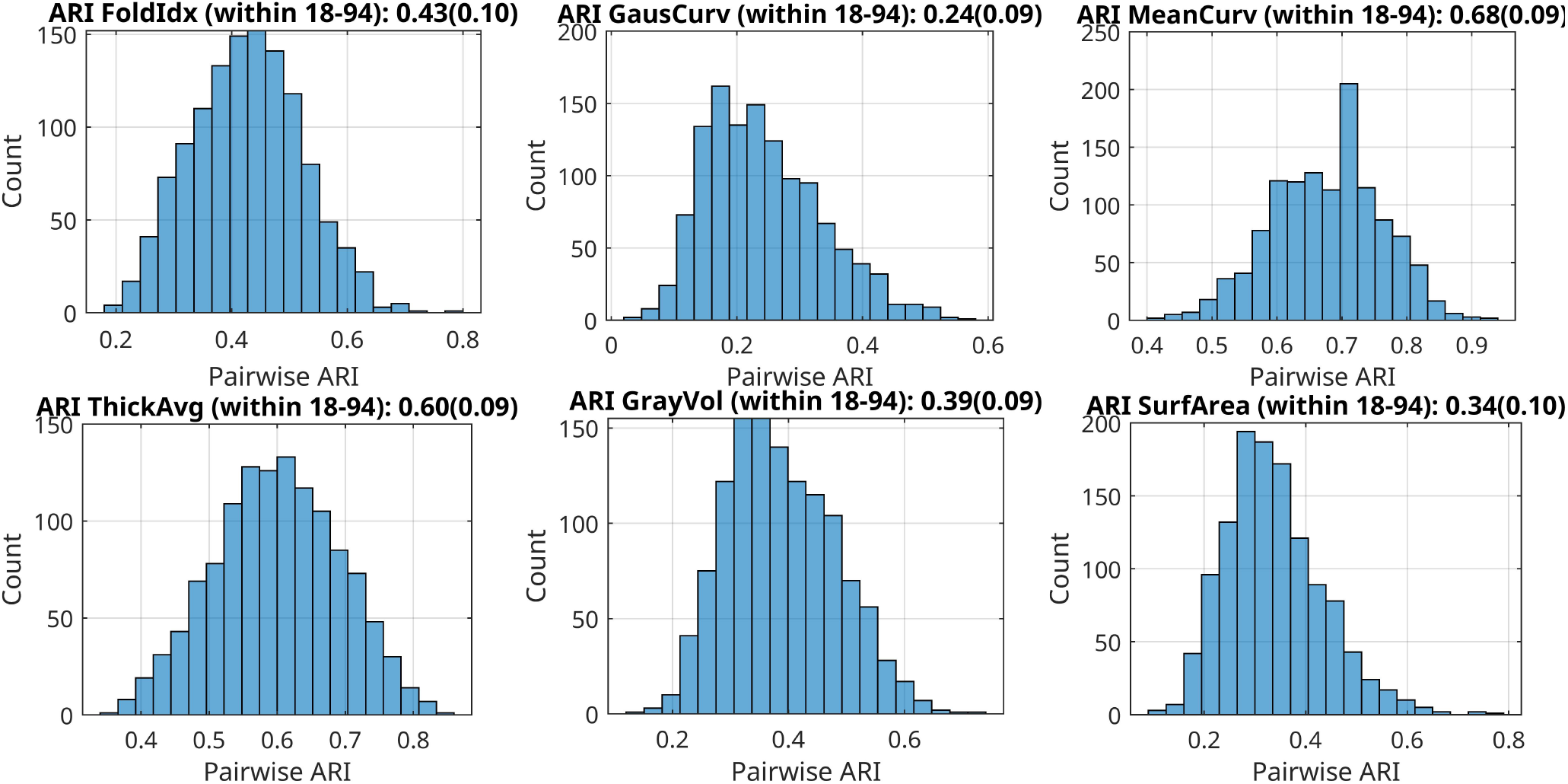
Feature-level within-age-bin structural covariance network reorganisation. Distribution of ARIs within lifespan (18 – 94 years of age). Top panels: Curvature-based features; Bottom panels: Volume-based features. Abbreviation: Adjusted Rand index: ARI.

